# Association between quinolone use in food animals and gonococcal resistance to ciprofloxacin: an ecological study

**DOI:** 10.1101/2021.06.03.446933

**Authors:** Natalia Gonzalez, Said Abdellati, Sheeba Manoharan-Basil, Chris Kenyon

## Abstract

**Background:** Concentrations of fluoroquinolones up to 200-fold lower than the MIC have been shown to be able to select for antimicrobial resistance in E. coli and Salmonella spp. (the minimum selection concentration – MSC). We aimed to i) establish what the ciprofloxacin MSC is for Neisseria gonorrhoeae and ii) Assess at a country level if the prevalence of gonococcal ciprofloxacin resistance is associated with the concentration of quinolones used in food animal production (an important determinant of long-term low dose ciprofloxacin exposures in humans).

**Methods:** i). To assess if sub-inhibitory ciprofloxacin concentrations could select for de novo generated resistant mutants, susceptible WHO-P was serially passaged at 1, 1/10, 1/100 and 1/1000 of the ciprofloxacin MIC of WHO-P (0.004mg/L) on GC agar plates. ii) Spearman’s correlation was used to assess the association between the prevalence of ciprofloxacin resistance in N. gonorrhoeae and the two independent variables – quinolone use for animals and quinolone consumption by humans.

**Results:** Ciprofloxacin concentrations as low as 1/1000 of the MIC of WHO-P were able to select for ciprofloxacin resistance. The prevalence of ciprofloxacin resistance in *N. gonorrhoeae* was positively associated with quinolone use for food animals (ρ=0.47; P=0.004; N=34).

**Conclusion:** Further individual level research is required to assess if low doses of ciprofloxacin from ingested foodstuffs are able to select for ciprofloxacin resistance in N. gonorrhoeae and other species.

## Introduction

Why has gonococcal resistance to fluoroquinolones emerged so explosively in East Asia? The prevalence of ciprofloxacin resistance in China, for example, increased from 10% to 95% between 1996 and 2003 [1]. In comparison, the country-level median prevalence of ciprofloxacin resistance in 2009 was 24% in the Americas and 6% in Africa [2]. Whilst differences in the intensity of exposure to fluoroquinolones likely play a key role in explaining these differences in resistance, it is not clear what pathways are most important. Quinolones used to treat *N. gonorrhoeae* can select for antimicrobial resistance (AMR) directly (direct selection). Because *N. gonorrhoeae* is asymptomatic for the majority of the time it circulates in a population, quinolones used for other indications can also select for AMR (bystander selection) [3]. *N. gonorrhoeae* can also acquire resistance conferring genes from commensal *Neisseria* [4]. Quinolone used for any indication could select for ciprofloxacin resistance in commensals and subsequently in *N. gonorrhoeae* via this indirect selection pathway [5, 6]. Studies have provided support for each of these pathways in the genesis of gonococcal AMR [4-6]. In particular, a number of studies have established that country level consumption of quinolones is a predictor of ciprofloxacin resistance [2, 3]. Certain Asian countries such as China, Malaysia and the Philippines have been noted to be outliers in these analyses [2]. In 2009, for example, these 3 countries had close to 100% resistance to ciprofloxacin, but all consumed less than half the volume of quinolones consumed by the United States where less than 10% of gonococcal isolates were resistant to ciprofloxacin [2].

There is also an increasing body of evidence to suggest that quinolone resistance in *N. gonorrhoeae* in Asia and elsewhere is part of a syndemic of resistance affecting a range of gram negatives including *Escherichia coli* and *Pseudomonas spp*. [7]. Further evidence for the syndemic perspective comes from studies which show that the prevalence of quinolone resistance is considerably higher (up to 16-fold higher) in a number of gram negatives in Asia than in Europe and the Americas [8-11]. Between 1998 and 2009, for example the prevalence of ciprofloxacin resistance in *Shigella* increased from 0% to 29% in Asia compared to 0% to 0.6% in Europe-America [8]. This syndemic of quinolone resistance may include *N. meningitidis* and various species of commensal *Neisseria* [12]. Recent studies from Shanghai, for example, have found the prevalence of ciprofloxacin resistance to be 99% in *N. gonorrhoeae*, 100% in commensal *Neisseria* and 66% in *N. meningitidis* [6, 13, 14]. These findings suggest that quinolone exposure other than that used for human infections, may be playing a role in the genesis and spread of quinolone resistance. One such possibility is quinolones used for animal husbandry.

Quinolone use in animals has been linked to AMR in a number of gram-negative pathogens circulating in humans [15, 16]. This use of quinolones could induce resistance in *N. gonorrhoeae* directly or indirectly. Direct selection on *N. gonorrhoeae* would occur via human ingestion of quinolone residues in meat or water/soil contaminated by animal manure [17]. Quinolones have been found to show very low biodegradability in the environment [17, 18]. Selection could also occur indirectly where quinolones select for resistance in commensal *Neisseria* which is then transformed into *N. gonorrhoeae*. Commensal *Neisseria* have been found in the resident microbiomes of a range of food animals including chickens, cows, sheep and goats [19-22]. The selection of quinolone resistance in commensal *Neisseria* could thus occur in the animals or humans. Of note the indirect pathway has been shown to be important in the genesis and spread of cephalosporin resistance (mainly via the spread of plasmids) in various gram-negative bacteria such as *E. coli* [15].

An important finding in this field is that antimicrobial concentrations up to 230-fold lower than the minimal inhibitory concentration (MIC) are capable of inducing AMR in bacteria such as *E. coli* and *Salmonella enterica spp*. [23, 24]. Whilst such experiments have not been conducted in *Neisseria spp*., concentrations of ciprofloxacin as low as 0.1 μg/L have been shown to able to select for resistance in other gram negative bacteria [23, 25]. Quinolone concentrations in meat, water and environmental samples have been found to far exceed this threshold in a number of locales. For example, studies have found that the mean concentration of ciprofloxacin in samples of milk, eggs, and edible fish in China to be 8.5 µg/L, 16.8 µg/kg and 331.7 µg/kg respectively (Fig. 1)[26-28]. Not even drinking water from taps (median quinolone concentration 0.270 μg/L) [29] and bottled water [30] in China is free of quinolones and other antimicrobials.

**Figure 1.**
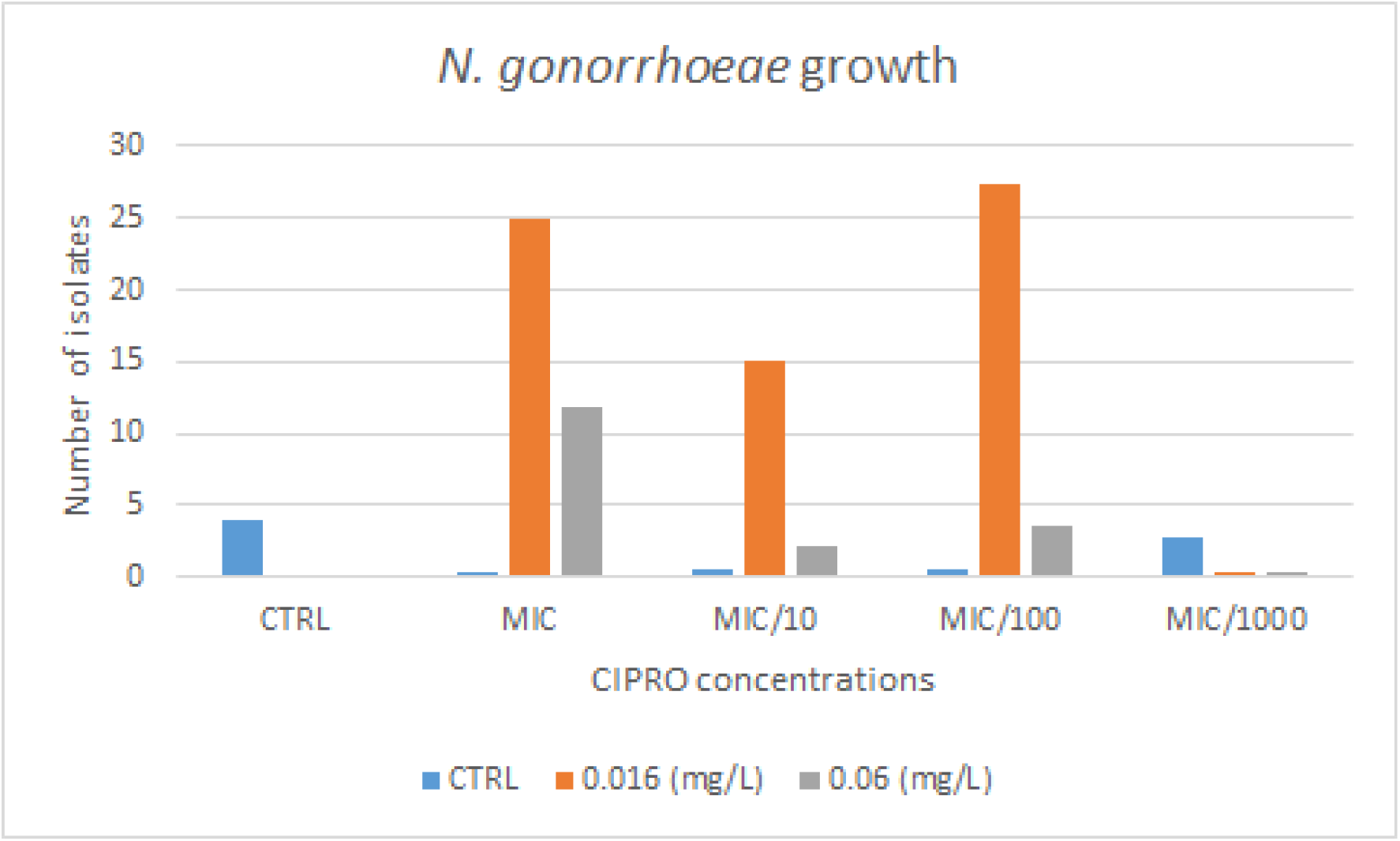
Mean number of colonies of *N. gonorrhoeae* grown for 7-days in different ciprofloxacin concentrations ranging from 1xMIC to 1/1000^th^ MIC of WHO-P, stratified according to their final growth in plates with no ciprofloxacin, 0.016 mg/L and 0.06mg/L. All the experiments were conducted in quadruplicate.

**Figure 2.**
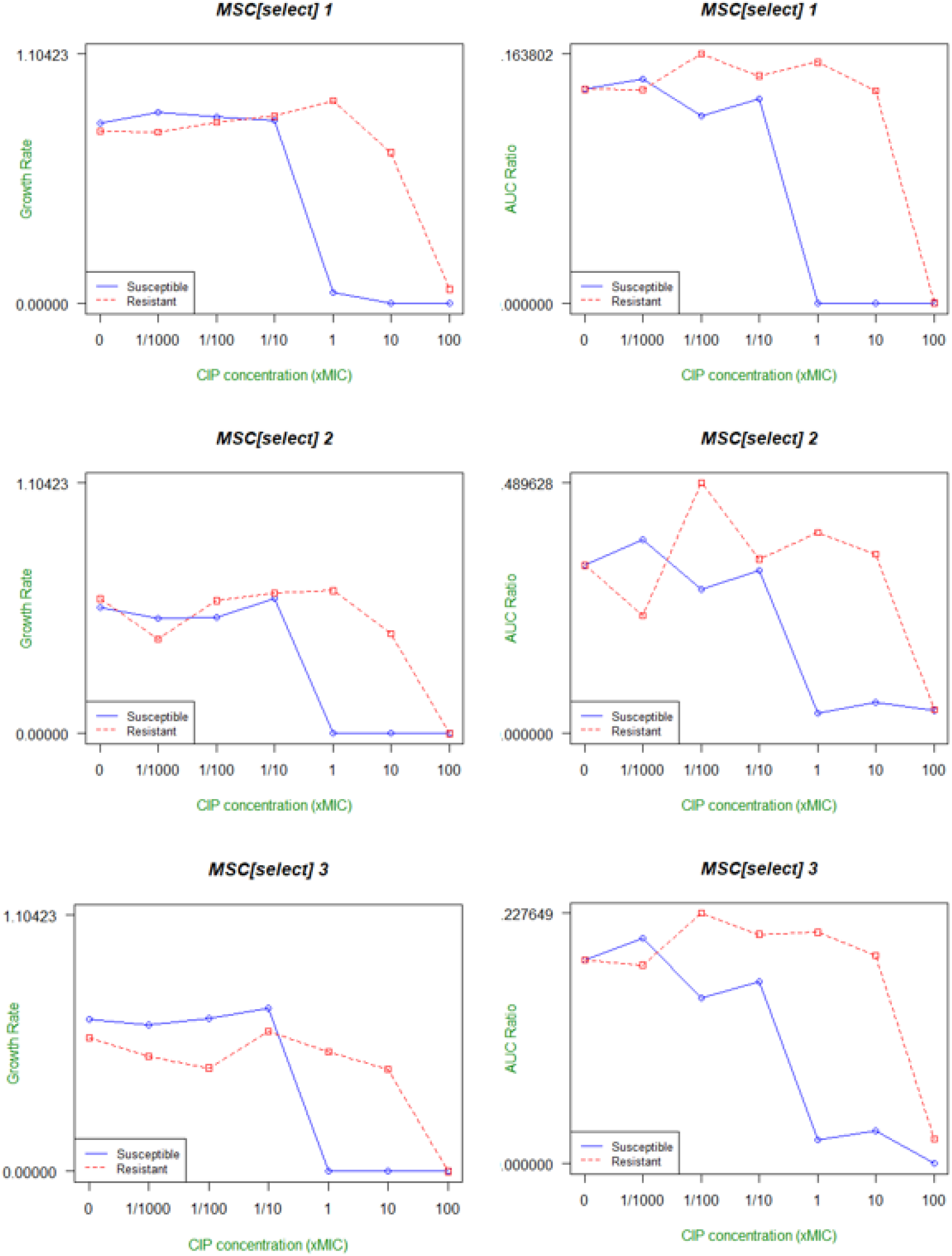
Growth rates as a function of ciprofloxacin (CIP) concentrations. The relative exponential growth rates (A, B and C) and AUC Ratio (D,E and F) of N. gonorrhoeae WHO-P with (red) and without (blue) the gyrA S91F mutation conferring resistance to ciprofloxacin. The MSCselect is determined as the point where the blue and red lines intersect.

These findings provided the motivation to for the two aims of this study: i) Assess the MSC of *N. gonorrhoeae*. ii) Assess at a country level if the prevalence of gonococcal ciprofloxacin resistance is associated with the concentration of quinolones used in food animal production.

## Methods

### A. In-vitro determination of minimum selection concentrations (MSC)

#### Bacterial strains and growth conditions

The WHO-P reference strain of *N. gonorrhoeae* was used for all experiments. The ciprofloxacin resistant strain was selected for by culturing on gonococcal (GC) agar plates supplemented with ciprofloxacin 0.008 mg/L (4 x MIC). After 24 hours the MIC of this strain attained 0.125 mg/L and whole genome sequencing confirmed the acquisition of the S91F resistance associated mutation in *gyrA*.

All experiments were conducted using GC broth supplemented with IsoVitaleX™ (1%) for liquid or solid media. Strains were grown at 36 °C, in 5% CO2.

#### MSC Determination

There are two important components of the MSC: i) The minimum concentration of an antimicrobial at which one can select for de-novo resistance (MSC_denovo_); ii) The lowest antimicrobial concentration that selects for a resistant-compared to a susceptible-strain (MSC_select_) [31, 32]. The methodology to assess the MSC_select_ and the MSC_denovo_ was based on that used by Gullberg et al [23, 25].

#### MSC_denovo_

To assess if sub-inhibitory ciprofloxacin concentrations could select for *de novo* generated resistant mutants, susceptible WHO-P was serially passaged at 1, 1/10, 1/100 and 1/1000 of the ciprofloxacin MIC of WHO-P (0.004mg/L) on GC agar plates. Identical control experiments were conducted with plates containing no ciprofloxacin. They were passaged at the same concentration of ciprofloxacin every 24 hours for 7 days. The number of colonies of WHO-P with reduced susceptibility and resistance to ciprofloxacin was established by plating 600 µL from a Phosphate Buffered Saline (PBS) solution containing the full growth from each experiment onto three GC agar plates: i) no ciprofloxacin, ii) ciprofloxacin concentration of 0.016 mg/L (reduced susceptibility) and iii) ciprofloxacin 0.06 mg/L (EUCAST breakpoint for ciprofloxacin resistance). The number of colonies was counted after 24 hours incubation. The lowest ciprofloxacin concentration that resulted in growth of WHO-P in the ciprofloxacin 0.06 mg/L plates was defined as the MSC_denovo._ Each experiment was conducted in quadruplicate. A subset of the colonies were replated on plates with the same concentration of ciprofloxacin to confirm that they were resistant.

#### MSC_select_

Growth rates were measured in standard 15ml vials in the NGmorbidostat whose construction and optimization for the culture of *N. gonorrhoeae* has been described elsewhere [33, 34]. The vials were continuously stirred at a speed of 200 rotations per minute (rpm) via a magnetic stirrer. The optic density (OD) of each vial was measured once a minute via a system of infrared light emitting diodes and photodetectors and recorded in an automated fashion (Matlab R2015b).

The growth rates of the susceptible and resistant *N. gonorrhoeae* at 36 °C in GC broth, were assessed in vials containing the following ciprofloxacin concentrations: 0, 1, 1/10, 1/100 and 1/1000 times of the ciprofloxacin MIC of WHO-P (0.004mg/L).

Each vial was inoculated with 100 µL of 4 McFarland *N. gonorrhoeae*, and the cultures were grown for 24 hours with continuous stirring. The relative growth rates at each concentration of ciprofloxacin were calculated as the observed growth rate of the strain divided by the growth rate of the same strain without ciprofloxacin. This experiment was conducted in triplicate.

Growthcurver was used to summarize the growth characteristics of each vial [35]. Growthcurver fits a basic form of the logistic equation common in ecology and evolution to experimentally-obtained growth curve data. It provides a number of summary measures to describe growth curves. We used two of these to calculate the MSC_select_

1. *Growth rate*. The intrinsic growth rate of the population, *r*, is the growth rate that would occur if there were no restrictions imposed on growth. Growthcurver uses the non-linear least-squares Levenberg-Marquardt algorithm to determine *r*.
2. *Area under the curve*. Growthcurver calculates the area under the logistic curve (AUC). This integrates information from the carrying capacity, growth rate and the population size at time 0.

The MSC_select_ was calculated as the point where the relative growth curves of the ciprofloxacin resistant and susceptible *Neisseria gonorrhoeae* crossed [25]. For the calculation of the MSC_select_, we calculated the average value obtained from the individual experiments.

### B. Ecological association between quinolone use and ciprofloxacin MICs

#### Quinolone use for animal food production data

We obtained the country level consumption of quinolones for animal food production in the year 2013 from a systematic review on this topic performed by Broeckel et al. [36]. This study calculated the volume of antimicrobials (in tons) by class of antimicrobial in 38 countries in the year 2013. Four categories of animals were included: chicken, cattle, pigs and small ruminants (sheep and goats), which together account for the overwhelming majority of terrestrial animals raised for food [15, 36].

We used this data to calculate the number of milligrams of quinolones used for animal food production/population correction unit (PCU) (a kilogram of animal product) in the year 2013. The data for the tonnage of food animals produced per country and year in the year 2013 was taken from the Food and Agriculture Organization estimates (http://www.fao.org/faostat/en/?#data/).

#### Quinolone consumption in humans

Data from IQVIA were used as a measure of national antimicrobial drug consumption. IQVIA uses national sample surveys that are performed by pharmaceutical sales distribution channels to estimate antimicrobial consumption from the volume of antibiotics sold in retail and hospital pharmacies. The sales estimates from this sample are projected with use of an algorithm developed by IQVIA to approximate total volumes for sales and consumption. Quinolone consumption (moxifloxacin, ciprofloxacin, gemifloxacin, ofloxacin, levofloxacin, lomefloxacin, norfloxacin, enoxacin, gatifloxacin, trovafloxacin, sparfloxacin) estimates are reported as the number of standard doses (a dose is classified as a pill, capsule, or ampoule) per 1000 population per year [2].

#### *N. gonorrhoeae* ciprofloxacin resistance data

The percent of isolates per country that were resistant to ciprofloxacin in the year 2014 (the year following the quinolone consumption variables) was taken from the WHO Global Gonococcal Antimicrobial Surveillance Programme (GASP; https://www.who.int/data/gho/data/indicators). GASP is a collaborative global network of regional and subregional reference laboratories that monitors gonococcal AMR in participating countries. The full GASP methodology, including suggested sampling strategy, laboratory techniques, external quality assurance, and internal quality control mechanisms has been published elsewhere [37]. GASP uses a minimum inhibitory concentration (MIC) breakpoint of 1μg/ml to define resistance to ciprofloxacin, which was therefore the definition of ciprofloxacin resistance we used [37]. In the case of 4 countries, data was not available for the year 2014. We used the data for the first subsequent year with available data – 2016 (Czechia, Luxembourg) and 2017 (Finland, Switzerland).

### Statistical analysis

Spearman’s correlation was used to assess the association between the prevalence of ciprofloxacin resistance in *N. gonorrhoeae* and the two independent variables – quinolone use for animals and quinolone consumption by humans. Linear regression was used to assess the country-level association between the percent of *N. gonorrhoeae* isolates with ciprofloxacin resistance and the two independent variables in 3 models. We started by assessing the association between ciprofloxacin resistance and quinolone consumption in humans (Model-1). We then assessed the association between ciprofloxacin resistance and quinolone use in animals (Model-2). Finally in Model-3 we evaluated the effect of both independent variables on ciprofloxacin resistance. Stata 16.0 was used for all analyses. A p-value of <0.05 was considered statistically significant.

## Results

### Minimum selection concentration

#### MSC_denovo_

Ciprofloxacin concentrations as low as 0.004μg/L (1/1000^th^ the MIC) were able to induce resistance to ciprofloxacin (the minimum selection concentration; Fig. 1; Supplementary Table1). This occurred in only one colony in one of four experiments at 0.004μg/L. Higher concentrations of ciprofloxacin resulted in a higher number of ciprofloxacin resistant colonies. Unsurprisingly, the number of colonies with reduced susceptibility to ciprofloxacin was generally higher than the number with ciprofloxacin resistance.

#### MSC_select_

*The mean MSC*_*select*_ *obtained by the AUC ratio method (mean 0*.*007* mg/L, *range 0*.*001-0*.*019*mg/L) *was considerably lower than that obtained by the growth rate method (mean 0*.*413* mg/L, *range 0*.*023-0*.*98*mg/L; Fig.1; Supplementary Table 2).

### Ecological association between quinolone consumption and ciprofloxacin resistance

No quinolone consumption data was available for two countries (Iran and Nepal). There was considerable variation in quinolone use for food animals in the 36 countries with data (median 1.9 mg quinolones/PCU (IQR 0.7-6.6 mg/PCU; Supplementary Table 2). Quinolone exposure in China was higher than all other countries (261.2 mg/PCU).

The prevalence of ciprofloxacin resistance in *N. gonorrhoeae* was positively associated with quinolone use for food animals (ρ=0.47; P=0.004; N=34) but not quinolone consumption in humans (ρ=0.31; P=0.097; N=30).

The model that combined quinolone use in food and humans was a better predictor of gonococcal ciprofloxacin resistance (R^2^ =0.30; Model 3) than the models that only included either quinolone use in animals or humans (R^2^=0.14 [Model 2] and R^2^=0.07 [Model 1], respectively; Table 2).

## Discussion

We found that the gonococcal MSC for ciprofloxacin was lower than that for any other bacteria on record [23, 25]. In particular the MSC_**denovo**_ (0.004μg/L) was considerably lower than quinolone concentrations found in foodstuffs such as meat products, milk and water in China and elsewhere (Fig. 3) [26-30]. The consumption of foods with high quinolone concentrations has been found to be associated with high urinary and faecal concentrations of quinolones in humans [38-41]. For example, a study from South Korea found that high urinary excretion of enrofloxacin and ciprofloxacin in the general population were strongly associated with consumption of beef, chicken and dairy products [38]. Likewise, a large study of the general population in 3 regions of China found ciprofloxacin, enrofloxacin and ofloxacin in the faeces of 67%, 30% and 57% of individuals [41]. The authors attributed the high median concentration of quinolones (median 20 μg/kg), in large part, to the ingestion of veterinary antimicrobials in food [42]. Reducing consumption of these foodstuffs has also been found to result in a reduction of urinary quinolone concentrations [42]. The problem of high antimicrobial concentrations in human food is not limited to Asian countries. In European countries, food animals consume a greater mass of antimicrobials per kilogram per year than humans do [43]. Furthermore, studies from Europe have shown AMC in animals is independently associated with AMR in bacterial species colonizing and infecting humans [43].

These findings generate the hypothesis that quinolone concentrations in food may play a role in the genesis of quinolone resistance in bacteria such as *N. gonorrhoeae*. We found supportive evidence at a country level – quinolone consumption for animal husbandry was positively associated with the prevalence of ciprofloxacin resistance in *N. gonorrhoeae*. Furthermore, the combination of quinolone consumption in humans and food animals provided the best prediction of ciprofloxacin resistance. These findings could explain a number of anomalies that have been noted in the global epidemiology of quinolone resistance in *N. gonorrhoeae* and other *Neisseria*. China has noted to be an outlier in the generally positive associations that have been found between country-level quinolone consumption in humans and resistance [2]. Quinolone consumption by humans is not in the top quintile but the prevalence of quinolone resistance in *N. gonorrhoeae* as well as *N. meningitidis* and commensal *Neisseria* is one of the highest in the world [2, 6, 37, 44].

Our estimates of the MSC_select_ derived from the two methods differed considerably. The estimate of the MSC_select_ derived from the AUC ratio method was however similar to the MSC_denovo_. A number of factors have been shown experimentally to affect the MSC. For example, MSCs have been shown to increase by 13-to 43-fold in complex environments such as those in the human body where bacteria are competing with one another and interacting with the hosts defenses [24, 45, 46]. The use of *gfp* labelling of the resistant and susceptible strains in direct competitive growth assays has also been shown to provide a more accurate way to ascertain the MSCselect [25].

Furthermore, in a real-world setting, human microbiota would be exposed to low dose antimicrobials intermittently (such as during and after eating) rather than continuously such as in our experiments. There are also large differences in antimicrobial penetration to different anatomical sites such as the oropharynx which we have not considered [47]. We also do not know if the effect of quinolones in food on *Neisseria* would act via direct contact with *Neisseria* in the mouth during mastication, in the colon during drug elimination or in the genital tract following absorption and distribution. We have also not considered the indirect-commensal-pathway through which quinolones could select for AMR. Quinolones could select for resistance in commensals which could then be transferred to the pathogenic *Neisseria* via transformation [6, 48]. These considerations imply that our estimates of the gonococcal MSC should be viewed as tentative. It is possible and even likely that the lowest concentration that can select for quinolone resistance in *Neisseria* species is contingent on a large number of cofactors such as microbiome community state types and individual human pharmacogenomic variations [24, 45, 46, 49].

For these reasons, we consider the calculation of an exact gonococcal ciprofloxacin MSC less important than establishing that it is considerably lower than the MIC. What is important is conducting experiments to ascertain if the concentrations of quinolones detected in contemporary foodstuffs can induce AMR in *N. gonorrhoeae* and other bacteria. Whilst it may to be difficult to do human challenge studies with *N. gonorrhoeae*, studies with commensal *Neisseria* may be possible. A further option would be to assess if food spiked with low concentrations of quinolones could induce AMR in *Neisseria musculi*, a colonic and oropharyngeal commensal of the common mouse, *Mus musculus* [50, 51]. These studies may be of considerable use in determining what safe maximum residue limits (MRLs) of quinolones are in foodstuffs. Current European Commission and WHO/FAO guidelines for establishing MRLs evaluate the effect of antimicrobial residues on the toxicity towards a range of bacteria [52, 53]. They do not however evaluate the effect of these residues on the genesis of AMR [46, 52, 54, 55].

There are also a number of limitations pertaining to the ecological study. These include the relatively small number of countries with available data, the lack of longitudinal data on quinolone consumption in animals and the absence of data on quinolone use for aquaculture. The epidemiology of resistance is complex and factors other than the amount of quinolones consumed may influence the level of quinolone resistance. These include consumption of other classes of antimicrobials, travel by humans and trade of live animals and meat [15].

Our results are thus best considered hypothesis generating. Further in-vivo experiments along the lines outlined above will be required to assess if quinolones in food residues are playing a role in the genesis of quinolone resistance in *N. gonorrhoeae* and related bacteria.

## Authors’ contributions

NG and CK conceptualized the study. NG and SA conducted the MSC experiments and CK was responsible for the acquisition, analysis and interpretation of the ecological analyses. All authors read and approved the final draft.

## Competing interests

The authors declare that they have no competing interests.

## Funding

Nil

## Acknowledgements

Nil

## Consent for publication

Not applicable

**Table 1.** Linear regression models testing the country-level association between quinolone consumption in food animals/humans and the prevalence ciprofloxacin resistance expressed as a percentage [coefficients (95% confidence intervals)]

|  | Model 1 | Model 2 | Model 3 |
| --- | --- | --- | --- |
| Quinolones Humans | 0.02 (-0.007-0.043) | - | 0.2 (0.001-0.046)* |
| Quinolones Food animals | - | 0.2 (0.02-0.38)* | 0.2 (0.07-0.40)** |
| N | 30 | 34 | 30 |
| R<sup>2</sup> | 0.07 | 0.14 | 0.30 |

\* P-value <0.05, \*\* P-value <0.01

## Notes

### Competing Interest Statement

The authors have declared no competing interest.

### Summary of Updates

Updated version including MSC experiments

