## Supplementary FIle for "Association between quinolone use in food animals and gonococcal resistance to ciprofloxacin: an ecological study"

Supplementary Table 1. Quinolone consumption in humans (defined daily doses per 1000 inhabitants per year) and in food animals (mg/population correction unit (PCU)) and prevalence of ciprofloxacin resistance in *N. gonorrhoeae*

| Country | Quinolone consumption humans, 2013 | *N. gonorrhoeae* ciprofloxacin resistance (%), 2014 | Quinolone use in food animals mg/PCU, 2013 |
| --- | --- | --- | --- |
| Australia | 264 | 36.43 | 0 |
| New Zealand | NA | 47.34 | 0.0305 |
| Sweden | 369 | 57 | 0.0792 |
| Canada | 874 | 34.02 | 0.085 |
| Iceland |  | 58.33 | 0.123 |
| Finland | 452 | 42 | 0.2495 |
| USA | 1099 | 19.2 | 0.402 |
| United Kingdom | 253 | 33.33 | 0.6592 |
| Austria | 684 | 52.48 | 0.6641 |
| Switzerland | 790 | 40.5 | 0.8424 |
| Denmark | 311 | 34.86 | 0.8955 |
| Ireland | 432 | 34.65 | 0.9568 |
| Netherlands | 374 | 32.16 | 1.037 |
| Germany | 729 | 63.21 | 1.4234 |
| Cyprus | NA | 100 | 1.5842 |
| France | 770 | 50.91 | 1.6459 |
| Japan | 999 | 76.05 | 1.6865 |
| Belgium | 1173 | 57.86 | 1.8031 |
| Norway | 304 | 73.64 | 2.036 |
| Lithuania | 467 | NA | 2.2816 |
| Czech Republic | 396 | 52.2 | 2.4273 |
| Slovenia | 504 | 45.12 | 2.5138 |
| Vietnam | 732 | 98.67 | 2.542 |
| Estonia | 368 | 15.38 | 2.8088 |
| Luxembourg | 1123 | 70 | 3.2984 |
| Latvia | 411 | 19.05 | 4.9641 |
| Slovakia | 982 | 67.89 | 4.9681 |
| Italy | 1520 | 78 | 8.1678 |
| Poland | 574 | 65.22 | 8.8151 |
| Hungary | 907 | 55.06 | 8.873 |
| Portugal | 1011 | 36.36 | 10.2723 |
| Spain | 1151 | 67.55 | 12.4847 |
| Bulgaria | 1057 | NA | 13.4787 |
| Sri Lanka | NA | 99.06 | 20.4333 |
| South Korea | 717 | 95 | 23.5592 |
| China | 316 | 99.6 | 261.152 |

NA- not available

Supplementary Table 2. Number of colonies of *N. gonorrhoeae* grown in plates at 0.06, 0.016 and 0 (mg/L) of ciprofloxacin after 7 days of subculturing exposed to different concentrations of it.

|  | CTRL | 0,016 (µg/mL) | 0,06 (µg/mL) |
| --- | --- | --- | --- |
| 1.CTRL | 3 | 0 | 0 |
| 2.CTRL | 3 | 0 | 0 |
| 3.CTRL | 8 | 0 | 0 |
| 4.CTRL | 2 | 0 | 0 |
| 1.MIC1 | 1 | 0 | 0 |
| 2.MIC1 | 0 | 0 | 6 |
| 3.MIC1 | 0 | >50 | 1 |
| 4.MIC1 | 0 | >50 | 40 |
| 1.MIC/10 | 0 | 2 | 1 |
| 2.MIC/10 | 2 | 8 | 2 |
| 3.MIC/10 | 0 | 0 | 0 |
| 4.MIC/10 | 0 | >50 | 6 |
| 1.MIC/100 | 2 | 8 | 1 |
| 2.MIC/100 | 0 | 1 | 0 |
| 3.MIC/100 | 0 | >50 | 5 |
| 4.MIC/100 | 0 | >50 | 8 |
| 1.MIC/1000 | 11 | 0 | 0 |
| 2.MIC/1000 | 0 | 0 | 0 |
| 3.MIC/1000 | 0 | 0 | 0 |
| 4.MIC/1000 | 0 | 1 | 1 |

Supplementary Table 3. The MSC_select_ values for N. gonorrhoeae WHO-P as derived from assessement of differential growth rates and AUC ratios.

| Experiment number | Growth Rate | AUC Ratio |
| --- | --- | --- |
| MSC1 (mg/L) | 0.238 | 0.001 |
| MSC2 (mg/L) | 0.023 | 0.019 |
| MSC3 (mg/L) | 0.98 | 0.0126 |
| Mean Result | 0.413 | 0.007 |
